# HiPhase: Jointly phasing small and structural variants from HiFi sequencing

**DOI:** 10.1101/2023.05.03.539241

**Authors:** James M. Holt, Christopher T. Saunders, William J. Rowell, Zev Kronenberg, Aaron M. Wenger, Michael Eberle

## Abstract

**Background:** In diploid organisms, phasing is the problem of assigning the heterozygous variants to one of two haplotypes. Reads from PacBio HiFi sequencing provide long, accurate observations that can be used as the basis for both calling and phasing variants. HiFi reads also excel at calling larger classes of variation such as structural variants. However, current phasing tools typically only phase small variants, leaving larger structural variants unphased.

**Methods:** We developed HiPhase, a tool that jointly phases SNVs, indels, and structural variants. The main benefits of HiPhase are 1) dual mode allele assignment for detecting structural variants, 2) a novel application of the A*-algorithm to phasing, and 3) logic allowing phase blocks to span breaks caused by alignment issues around reference gaps and homozygous deletions.

**Results:** In our assessment, HiPhase produced an average phase block NG50 of 493 kb with 933 switchflip errors and fully phased 95.2% of genes, improving over the current state of the art. Additionally, HiPhase jointly phases SNVs, indels, and structural variants and includes innate multi-threading, statistics gathering, and concurrent phased alignment output generation.

**Availability:** https://github.com/PacificBiosciences/HiPhase

## 1 Introduction

For human genomics, phasing variants within a gene is a critical aspect of clinical testing for pharmacogenomics (Scott et al., 2022; Graansma et al., 2023; van der Lee et al., 2020; Caspar et al., 2021), HLA typing (Mayor et al., 2019), and autosomal recessive conditions (Tewhey et al., 2011). Additionally, automated systems can benefit from knowing whether two variants are in *cis* or *trans* within a gene, potentially reducing the time for clinical analysts to reach a molecular diagnosis (Danecek and McCarthy, 2017; Owen et al., 2022). Beyond clinical diagnostics, there are many additional applications of phasing in genomics, many of which are summarized in a review by Browning and Browning (2011).

Haplotype phasing generally falls into two categories: statistical phasing (Browning and Browning, 2011) and read-backed phasing (Snyder et al., 2015). For read-backed phasing, each read is mapped to a reference genome and variants are called from the read mappings. The heterozygous variant calls are phased into a single diplotype composed of two haplotypes. Once phased, the variants can be fed into various tertiary analysis tools for downstream analyses. In practice, reads that are long and accurate, such as PacBio HiFi reads (Wenger et al., 2019), have a large number of informative bases overlapping heterozygous variants and are ideal for variant calling followed by read-backed phasing.

For HiFi reads, WhatsHap (Patterson et al., 2015) is the most prominent read-backed phaser and the *de facto* standard in the field for phasing single-nucleotide variants (SNVs) and short indels. WhatsHap uses a “fixed parameter tractable approach”, meaning that it down-samples the sequencing reads to a fixed coverage value, to phase the heterozygous variants (Patterson et al., 2015). Despite being generally efficient and accurate for phasing a human genome, WhatsHap does have some notable limitations: 1) it down-samples the data in order to run efficiently, potentially biasing the data; 2) multi-allelic variants (sites with two or more non-reference alleles) are not phased; 3) it does not span gaps in coverage caused by large deletions or reference gaps; 4) it is single-threaded, so parallel processing requires some engineering overhead; and it is limited to small variants such as SNVs and short indels. Recently, some tools have focused on that last limitation by either overlaying structural variants (insertions and deletions *≥* 50 bp) on the WhatsHap phase blocks (Mahmoud et al., 2021) or jointly phasing SNVs and structural variants, but not short indels (Lin et al., 2022). This creates a notable gap for a tool that accurately and jointly phases SNVs, indels, and structural variants.

Here we introduce HiPhase, a new tool that jointly phases SNVs, indels, and structural variants called from PacBio HiFi sequencing on diploid organisms. HiPhase uses two novel approaches to solve the phasing problem: 1) dual mode allele assignment and 2) a phasing algorithm based on the A* search algorithm (Hart et al., 1968). In addition to jointly phasing small variants with structural variants, HiPhase offers additional benefits such as: 1) no down-sampling of the data, 2) support for multi-allelic variation, 3) logic to span coverage gaps with supplementary alignments, 4) innate multi-threading, 5) built-in statistics gathering, and support for assigning aligned reads to a haplotype (“haplotagging”) while phasing. In our assessment, HiPhase produced an average phase block NG50 of 493 kb with 933 switchflip errors and fully phased 95.2% of genes.

## 2 Methods and Materials

HiPhase breaks the phasing problem into three major components: phase block generation, allele assignment, and diplotype solving.

### 2.1 Phase block generation

Phase block generation works by identifying pairs of consecutive heterozygous variant calls connected by at least one read mapping. A phase block is initialized by taking the first available variant on a chromosome and creating a single-variant block from it. Then, the next variant is loaded and HiPhase searches for mappings that span both that new variant and the current block. If no mappings span both, the algorithm will then check for supplementary mappings from the new variant into the current block. This supplementary mapping check allows HiPhase to span coverage gaps caused by homozygous deletions and reference gaps. If at least one spanning or supplementary mapping is identified, then the current phase block is extended to include the variant. Otherwise, the current block is returned as a putative phase block and a new single-variant block is initialized from the new variant. This process repeats until all heterozygous variants on the chromosome are consumed.

Each putative phase block acts as an isolated sub-problem in the full solution. Because the putative blocks are unconnected by the read mappings at their ends, they represent a lower-bound on the number of phase blocks in the final solution. Importantly, each sub-problem can be solved independently, allowing for allele assignment and diplotype solving to be performed in parallel for each putative phase block. This forms the basis for multi-threading in HiPhase.

### 2.2 Allele assignment

Allele assignment converts all read mappings within a putative phase block into condensed allelic observations which are chains of reference or alternate alleles corresponding to the observed alleles within a particular read mapping. This simplifies a long-read sequence down to a smaller set of integer values representing alleles that are present within the read. We refer to these assignments as reference or alternate alleles, but HiPhase also allows for ambiguity, unassigned alleles, and multi-allelic variation (two different alternate alleles at one position). Additionally, each observed allele is assigned a “weight” indicating the cost to alter or ignore that allele in the diplotype solving step.

HiPhase has two modes for allele assignment: local re-alignment and global re-alignment. In brief, local re-alignment assigns alleles and quality based on a small window around each variant position, which is conceptually similar to the allele assignment process of WhatsHap (Patterson et al., 2015). In contrast, global re-alignment will fully re-align the mapping against a local alt-aware reference graph using a graph-aware version of the wavefront algorithm (Marco-Sola et al., 2020). In general, local re-alignment is a faster process, but it is ill-suited for accurate allele assignment for large structural variants and even some smaller indels. Global re-alignment tends to be slower but is more accurate when it comes to allele assignment, especially around structural variants. Finally, HiPhase implements a “dual mode” allele assignment where if global re-alignment is too slow, it will fall back on local re-alignment for the phase block.

Once all mappings have been converted to a condensed allele representation, HiPhase collapses mappings with the same read name into a single entry. The primary purpose of this step is to create a bridge between supplementary mappings that span a gap in coverage. This allows HiPhase to cross deletion events and reference gaps bridged by split read mappings. If the mappings for one read overlap but have a conflicting allele assignment, then that allele is converted to an ambiguous allele assignment in the collapsed representation. In the end, each read is represented exactly once in the collection of condensed alleles for the phase block.

### 2.3 Diplotype Solving

Diplotype solving distills the condensed allelic observations into two representative haplotypes that are expected to complement each other (i.e., where one is the reference allele, the other is the alternate allele). For our purposes, we define the core diplotype solving problem as a slight reformulation of the weighted minimum error correction (wMEC) problem as described previously (Patterson et al., 2015). Given a matrix where each row corresponds to the condensed allele representation of one read in the phase block, the goal is to find two haplotypes, *h*_1_ and *h*_2_, such that the cost of changing each row to exactly match either *h*_1_ or *h*_2_ is minimized. We refer the reader to Patterson et al. (2015) for greater technical details around this and other formulations of the phasing / wMEC problem.

HiPhase uses a version of the A* search algorithm (Hart et al., 1968) to search for the optimal diplotype. In general, A* search algorithms explore a given search space by iteratively expanding the current lowest cost option based on both an observed cost and heuristic cost estimate. In HiPhase, the search space is all possible diplotype solutions modeled as a search tree, and the heuristic is generated by chaining sub-problem solutions. While increased coverage and variant density will increase the run-time of this algorithm, in practice we find that HiPhase can operate efficiently on the typical 30x human whole genome without down-sampling. Greater details on all HiPhase methods, heuristics, and limitations can be found in the Supplemental Materials.

### 2.4 Data

The following sections outline public resources used in our analysis. Details on and links to each can be found in the Supplemental Materials.

#### 2.4.1 Benchmark datasets

For benchmarking phase accuracy, we used publicly available Genome in a Bottle (GIAB) phased variant sets v4.2.1 (GRCh38) for HG001, HG002, and HG005 (Wagner et al., 2022). We note that these files are limited to the human autosomes, and do not provide phasing information for structural variants.

#### 2.4.2 HiFi sequencing datasets

We used publicly available HiFi sequencing datasets for evaluating HiPhase. Three replicates for the sample HG002 sequenced on the Revio system were obtained from https://downloads.pacbcloud.com/public/revio/2022Q4/. In our Supplemental Materials, we also compare separate datasets for HG001, HG002, and HG005 from the Sequel II system that were obtained from the GIAB consortium (Zook et al. (2016), https://ftp-trace.ncbi.nlm.nih.gov/ReferenceSamples/giab/data/).

#### 2.4.3 Pre-processing

We followed the recommended practices for pre-processing our HiFi datasets through variant calling. In brief, we aligned reads to GRCh38 using `pbmm2` and then performed variant calling with DeepVariant with the PacBio model (Poplin et al., 2018) and `pbsv`.

### 2.5 Phasing metrics

We measured the quantity of phasing errors with switchflips (Patterson et al., 2015). To measure phase block length, we calculated NG50 using GRCh38 for reference length. We additionally gathered the total number of phased variants, number of phased structural variants (HiPhase only), and the percentage of fully phased genes. We note that NG50 and number of phased variants are measured using tool-specific outputs. Further details on how all metrics are defined, calculated, and collected are available in the Supplemental Materials.

### 2.6 Tool descriptions

We focus our analyses on comparing the current approach for HiFi read-backed phasing, WhatsHap (Patterson et al., 2015), to our new tool, HiPhase. For WhatsHap, we provide small variants from DeepVariant and instruct the tool to phase both SNVs and indels. For HiPhase, we provide the same DeepVariant VCF file and an additional structural variant VCF file from `pbsv`. We set HiPhase to used global re-alignment (see Section 2.2) which is our recommended approach for phasing with structural variants. Additional results for other modes of running both WhatsHap and HiPhase are available in the Supplemental Materials.

## 3 Results

On average, HiPhase generated 493 kb phase block NG50, fully phased 95.2% of genes, and phased 3.1M variants with only 933 switchflip errors (see Figure 1), a rate of 1 error every 3.3K variants. Additionally, HiPhase generated over 25K phased structural variants per replicate, a feature that is not present in WhatsHap. Additional analyses with more metrics and samples are available in the Supplemental Materials.

**Figure 1:**
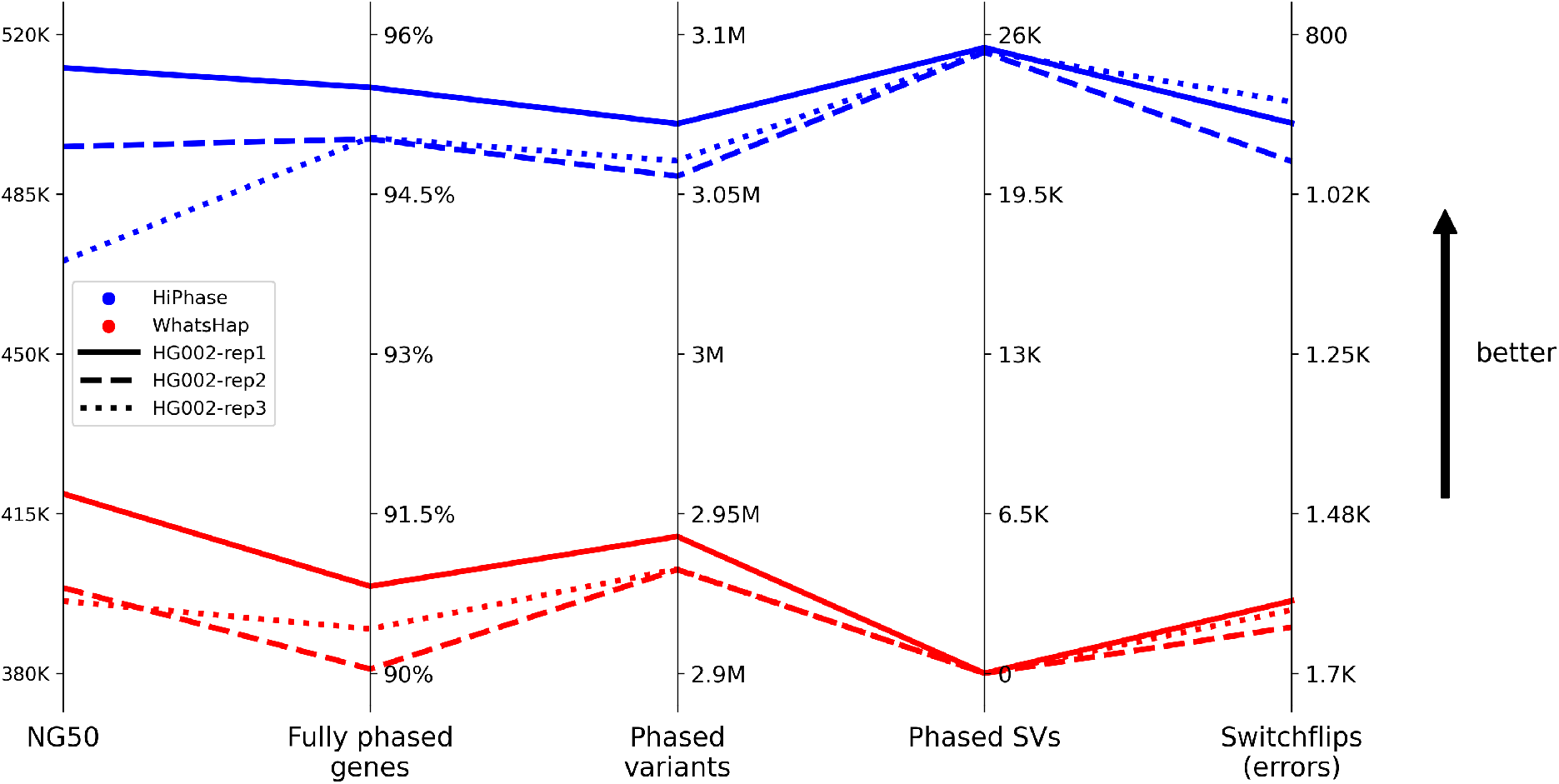
Summary comparison metrics. Each line represents a phase result where the line style corresponds to a HG002 replicate and the color corresponds to the phasing approach (blue for HiPhase or red for WhatsHap). Each metric axis is oriented such that better results are toward the top (i.e., switchflips is inverted). In all metrics, results from HiPhase were strictly better than those from WhatsHap.

To our knowledge, HiPhase is the first phasing tool to jointly phase SNVs, indels, and structural variants. Compared to the current approach, HiPhase generated longer phase blocks with fewer phasing errors, phased more total variants, and fully phased more genes in the human reference genome. HiPhase also includes innate multi-threading, statistics gathering, and concurrent phased alignment output generation. In future versions of the tool, we will extend HiPhase into other forms of structural variants such as inversions, tandem repeats, and copy number variation.

## 4 Availability

HiPhase is released as a pre-compiled binary file on GitHub: https://github.com/PacificBiosciences/ HiPhase. Additional instructions and use cases are available in the accompanying documentation. Details and links to all benchmark files, sequencing datasets, tool versions, and commands used in this document are available in the Supplemental Materials.

