## Supplemental Materials for "HiPhase: Jointly phasing small and structural variants from HiFi sequencing"

### Supplemental Material for "HiPhase: Jointly phasing small and structural variants from HiFi sequencing"

PacBio, Menlo Park, CA USA

3rd May, 2023

### 1 Data details

#### 1.1 Benchmark files

We benchmarked against the Genome in a Bottle (GIAB) consortium phased variant calls v4.2.1 (Wagner et al., 2022). Table 1 contains direct links to the files downloaded from GIAB for each of the three benchmark samples.

Table 1: Links to the phased benchmark set, GIAB v4.2.1, used for each sample.

| Sample | Benchmark file |
|---|---|
| HG001 | https://ftp-trace.ncbi.nlm.nih.gov/ReferenceSamples/giab/release/NA12878_HG001/NISTv4.2.1/GRCh38/SupplementaryFiles/HG001_GRCh38_1_22_v4.2.1_benchmark_hifiasm_v11_phasetransfer.vcf.gz |
| HG002 | https://ftp-trace.ncbi.nlm.nih.gov/ReferenceSamples/giab/release/AshkenazimTrio/HG002_NA24385_son/NISTv4.2.1/GRCh38/SupplementaryFiles/HG002_GRCh38_1_22_v4.2.1_benchmark_hifiasm_v11_phasetransfer.vcf.gz |
| HG005 | https://ftp-trace.ncbi.nlm.nih.gov/ReferenceSamples/giab/release/ChineseTrio/HG005_NA24631_son/NISTv4.2.1/GRCh38/SupplementaryFiles/HG005_GRCh38_1_22_v4.2.1_highconf_hifiasm_v11_phasetransfer.vcf.gz |

#### 1.2 Sequencing data

Links to the sequencing data used for all analyses can be found in Table 2. For HG002 and HG005 datasets from Sequel II systems, we used a subset of all available sequencing data to achieve approximately 30x read depth after alignment. All other datasets used every SMRT Cell that was available.

Table 2: Table with links to sequencing datasets used in this document. Datasets marked with an asterisk (*) used a subset of all available movies to reach approximately 30x. These datasets have the exact SMRT Cells used listed. All others are marked with "All" and the number of SMRT Cells available for use.

| System | Dataset | SMRT Cells used | URL |
|---|---|---|---|
| Sequel II | HG001 | All (6) | https://ftp-trace.ncbi.nlm.nih.gov/ReferenceSamples/giab/data/NA12878/HudsonAlpha_PacBio_CCS/ |
| | HG002* | m64012_190920_173625<br>m64012_190921_234837<br>m64015_190920_185703 | https://ftp-trace.ncbi.nlm.nih.gov/ReferenceSamples/giab/data/AshkenazimTrio/HG002_NA24385_son/PacBio_CCS_15kb_20kb_chemistry2/reads/ |
| | HG005* | m64017_200723_190224<br>m64109_200304_195708<br>m64109_200309_192110<br>m64109_200311_013444 | https://ftp-trace.ncbi.nlm.nih.gov/ReferenceSamples/giab/data/ChineseTrio/HG005_NA24631_son/HudsonAlpha_PacBio_CCS/ |
| Revio | HG002-rep1 | All (1) | https://downloads.pacbcloud.com/public/revio/2022Q4/HG002-rep1/ |
| | HG002-rep2 | All (1) | https://downloads.pacbcloud.com/public/revio/2022Q4/HG002-rep2/ |
| | HG002-rep3 | All (1) | https://downloads.pacbcloud.com/public/revio/2022Q4/HG002-rep3/ |

Summary statistics for each analyzed dataset are available in Table 3. Each dataset has approximately 30x sequencing depth. The three HG002 replicates sequenced on the Revio system have lower mean read lengths than HG001 or HG005, but they have a tail of longer reads as evidenced by higher N10 and N25

Table 3: Summary metrics for each dataset used in our analysis. Mean coverage was gathered after alignment from `mosdepth` (Pedersen and Quinlan, 2017a). Read length statistics were collected from `fastleng`.

| System | Dataset | Mean coverage | Mean read length | N25 read length | N10 read length |
|---|---|---|---|---|---|
| Sequel II | HG001 | 25.99x | 17,539 | 19,800 | 23,502 |
| | HG002 | 28.40x | 12,858 | 13,702 | 14,516 |
| | HG005 | 31.81x | 17,390 | 19,556 | 21,347 |
| Revio | HG002-rep1 | 32.38x | 15,474 | 20,696 | 24,342 |
| | HG002-rep2 | 29.59x | 15,296 | 20,465 | 24,027 |
| | HG002-rep3 | 27.96x | 15,247 | 20,407 | 24,072 |

read lengths. Compared to the other datasets, the HG002 dataset sequenced on the Sequel II system has noticeable shorter read lengths.

#### 1.3 Tool and pipeline versions

Table 4 contains the versions and links for all tools and pipelines that were used to generate results in this document.

#### 1.4 Tool definitions

The following list describes each of the tools used in our analysis, both in the main document and this supplement.

- WhatsHap - The current recommended approach for WhatsHap (Patterson et al., 2015) on HiFi datasets. It phases small variants from DeepVariant, including both SNVs and indels. This is the WhatsHap approach used for analyses in the primary document.
- WhatsHap (optimized) - Through experimentation, we found that allowing WhatsHap to distrust the genotypes tended to reduce the number of errors generated in the result at the cost of phase block length. However, this has the additional side-effect of allowing the tool to change heterozygous genotypes to homozygous in the output VCF files.
- HiPhase (no SV) - The method described in this paper when provided only the DeepVariant calls (i.e., no structural variants). This approach uses local re-alignment for allele assignment and is the recommend approach for phasing only small variants.
- HiPhase - This approach jointly phases DeepVariant calls with the large insertion and deletion calls from `pbsv`. This method uses global re-alignment and is our recommended approach for phasing with structural variants. This is the HiPhase approach used for analyses in the primary document.

The following sections contain the command templates used for each tool in our comparative analysis.

##### 1.4.1 WhatsHap

Program 1 contains the template we used for WhatsHap. For both WhatsHap variants, we parallelized by running one chromosome per job. This is specified in the {`wildcards.chrom`} parameter. The only other notable parameter is `--indel`, which enables the phasing of indels with SNVs.

##### 1.4.2 WhatsHap (optimized)

Program 2 is identical to the above, but with the added `---distrust-genotypes` option. This option will modify the algorithm to allow it to convert heterozygous calls into homozygous calls. We found that this typically reduces the number of errors generated by WhatsHap and reduces phase block length (NG50),

Table 4: Tool versions and URLs. This table contains primary tools used in this document as well as pipeline resources more upstream processing. Each tool or pipeline is tagged with the version used in this document and a link to the public repository.

| **Tool / method name** | **Version(s)** | **URL** |
|---|---|---|
| HiPhase (this paper) | v0.8.0 | https://github.com/PacificBiosciences/HiPhase |
| WhatsHap (Patterson et al., 2015) | v1.4 | https://github.com/whatshap/whatshap |
| Secondary Pipeline | – | https://github.com/PacificBiosciences/pb-human-wgs-workflow-snakemake |
| Reference genome | GRCh38 no alt analysis set | ftp://ftp.ncbi.nlm.nih.gov/genomes/all/GCA/000/001/405/GCA_000001405.15_GRCh38/seqs_for_alignment_pipelines.ucsc_ids/GCA_000001405.15_GRCh38_no_alt_analysis_set.fna.gz |
| pbmm2 | v1.4.0 | https://github.com/PacificBiosciences/pbmm2 |
| DeepVariant (Poplin et al., 2018) | v1.3.0 (Sequel II system)<br>v.1.5.0 (Revio system) | https://github.com/google/deepvariant |
| pbsv | v2.8.0 | https://github.com/PacificBiosciences/pbsv |
| mosdepth (Pedersen and Quinlan, 2017a) | v0.2.9 | https://github.com/brentp/mosdepth |
| fastleng | v0.2.0 | https://github.com/HudsonAlpha/rust-fastleng |

Program 1: Command line template for the baseline WhatsHap phasing method.

```
whatshap phase \
  --indel \
  --ignore-read-groups \
  --chromosome {wildcards.chrom} \
  --output {output.vcf} \
  --reference {input.reference} \
  {input.vcf} \
  {input.bams}
```

but this leads to an overall increase in corrected phase block length (NGC50). Additionally, it has the side effect of modifying the output variants to homozygous (0/0 or 1/1) instead of leaving them as unphased heterozygous calls.

Program 2: Command line template for the WhatsHap (optimized) phasing method.

```
whatshap phase \
  --indel \
  --distrust-genotypes \
  --ignore-read-groups \
  --chromosome {wildcards.chrom} \
  --output {output.vcf} \
  --reference {input.reference} \
  {input.vcf} \
  {input.bams}
```

##### 1.4.3 HiPhase (no SV)

The command in Program 3 is the baseline HiPhase method that is most comparable to WhatsHap because it only phases small variants. It is also the recommended approach if only a small variant call file is available. Each BAM is specified via the `--bam` parameter (multiple times if the data is stored in multiple BAM files). Multi-threading is enabled via `--threads` and multiple statistics files are also output while running (`--stats-file`, `--blocks-file`, and `--summary-file`). We note that similar to "WhatsHap (optimized)", HiPhase is allowed to convert heterozygous calls to homozygous if that leads to a more optimal solution. However, in contrast to WhatsHap, these variants are not converted in the output and are instead left as unphased heterozygous calls in the VCF.

Program 3: Command line template for the HiPhase (no SV) phasing method.

```
hiphase \
  --threads 16 \
  --reference {input.reference} \
  --bam {input.bam1} \
  ...
  --bam {input.bamN} \
  --vcf {input.vcf} \
  --output-vcf {output.vcf} \
  --stats-file {output.stats} \
  --blocks-file {output.blocks} \
  --summary-file {output.summary}
```

##### 1.4.4 HiPhase

Program 4 is the *recommended* way to use HiPhase for jointly phasing small variants with structural variants. The main changes relative to the above command are 1) the addition of a second input and output structural variant VCF and 2) the enabling of global re-alignment with the `--global-realignment-cputime` parameter.

### 2 Additional Results

In the main document, we focus on the latest sequencing data generated by the Revio system. However, the publicly available samples that also have a GIAB benchmark are limited to HG002. In these sections, we provide extended results for the Revio system datasets, and we also provide results for Sequel II system datasets that extend to HG001, HG002, and HG005 samples.

Program 4: Command line template for the HiPhase phasing method.

```
hiphase \
  --threads 16 \
  --global-realignment-cputime 300 \
  --reference {input.reference} \
  --bam {input.bam1} \
  ...
  --bam {input.bamN} \
  --vcf {input.vcf} \
  --output-vcf {output.vcf} \
  --vcf {input.pbsv_vcf} \
  --output-vcf {output.pbsv_vcf} \
  --stats-file {output.stats} \
  --blocks-file {output.blocks} \
  --summary-file {output.summary}
```

#### 2.1 All methods comparison

Figure 1 and Table 5 show the summary comparison for all tool definitions in this supplement (see Section 1.4) restricted to datasets from the Revio system. Compared to the current practice ("WhatsHap"), both HiPhase conditions generated fewer errors (switchflips) and longer corrected phase blocks (NGC50). Additionally, there is a small but noticeable reduction in switchflips and increase in NGC50 when structural variants are provided to HiPhase. We also note that these results demonstrate the potential benefit of the "WhatsHap (optimized)" mode relative to the current WhatsHap. Despite decreasing NG50, the positive effects of reducing switchflip errors leads to an overall improvement in NGC50.

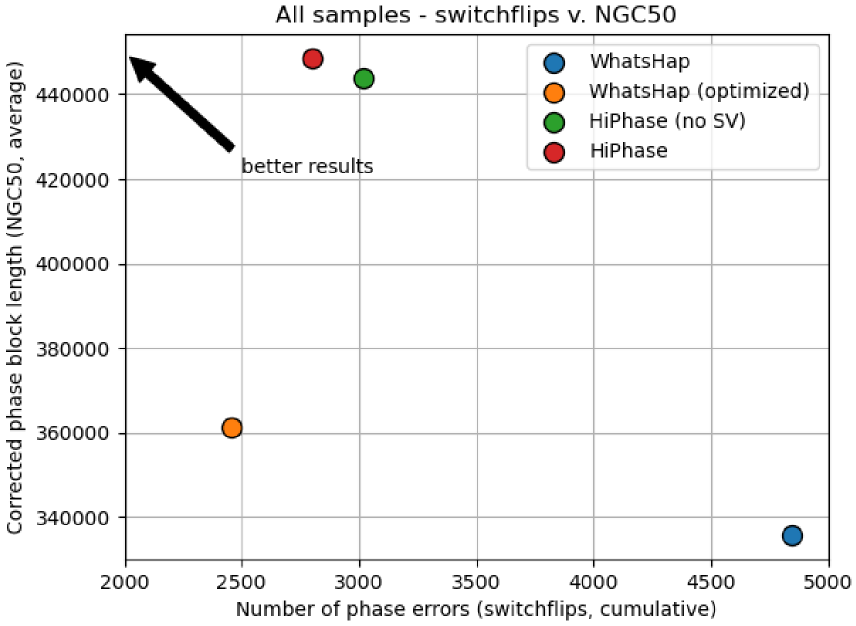

Figure 1: Summary figure showing switchflips v. NGC50 for all methods. Switchflips represent raw error counts whereas NG50 represents the length of error-free portions of the constructed phase blocks. HiPhase generated fewer switchflip errors than the primary comparator "WhatsHap", but more errors than the "WhatsHap (optimized)" version. However, NGC50 values were consistently higher for HiPhase than either WhatsHap method.

Table 5: Summary metrics for Revio system datasets (three replicates of HG002) for each method. Switchflips, hamming distance, and NGC50 are measures of error. NG50 and NGC50 are measures of phase block length. Number of phased variants is broken into "all" category and the structural variants (SVs) from `pbsv`. Switchflips, hamming distance, and the number of phased variants are totaled across all datasets (sum), whereas NG50, NGC50, and genes full phased are averaged (mean). Metrics marked with an asterisk (*) are derived from tool-specific sources, details of which are available in Section 3.1. **Bolded** values represent the best performance in the row. Similar to the results from the main document, HiPhase outperforms all other methods with the exception of switchflips and hamming distance from "WhatsHap (optimized)".

| Metric | WhatsHap | WhatsHap (optimized) | HiPhase (no SV) | HiPhase |
|---|---|---|---|---|
| Switchflips | 4,844 | **2,457** | 3,016 | 2,799 |
| Hamming distance | 148,201 | **91,356** | 166,184 | 142,505 |
| NG50* | 406,912 | 393,252 | 491,782 | **492,784** |
| NGC50 | 335,787 | 361,338 | 443,804 | **448,536** |
| Fully phased genes | 90.4% | 88.9% | 95.1% | **95.2%** |
| Number of phased variants (all)* | 8,807,866 | 8,652,027 | 9,128,275 | **9,188,036** |
| Number of phased SVs (`pbsv`) | N/A | N/A | N/A | **76,033** |

#### 2.2 Overall summary figure

Figure 2 shows the summary comparison for all datasets split by Sequel II or Revio systems and tool definitions. All methods on Revio system datasets produced higher NGC50 than the Sequel II system comparators. Additionally, HiPhase runs had a comparable switchflip counts, whereas WhatsHap tended to increase in error on Revio system datasets.

#### 2.3 Sequel II system summary metrics

Table 6 shows the more detailed summary results for Sequel II system datasets when evaluated on the four tool definitions. The patterns shown in these results generally reflect the same patterns observed in Revio system datasets.

### 3 Result metrics

All metrics were gathered using a benchmarking pipeline built with `snakemake` (Mölder et al., 2021). Where possible, metric gathering commands were homogenized. The following sections describe the overall summary of the metrics followed by specific commands as `snakemake` templates and highlight tool-based differences where appropriate.

#### 3.1 Metric definitions

##### 3.1.1 Error metrics

These metrics are generated when comparing a phase result to the GIAB benchmark sets:

- Switchflips - A "switch" is when a pair of consecutive heterozygous variants in the phase result are incorrectly phased relative to the benchmark set, and a "flip" is two back-to-back switches such that only one variant is on the incorrect phase block. "Switchflips" is the total number of switches and flips across all phase blocks (switches are not double-counted in a flip). Typically, this metrics is thought of as the *number* of errors contained in the phase result.

- Hamming distance - The minimum number of phase orientations that need to be "flipped" in the phase result to exactly match the benchmark set. This metric is heavily influenced by block length, number

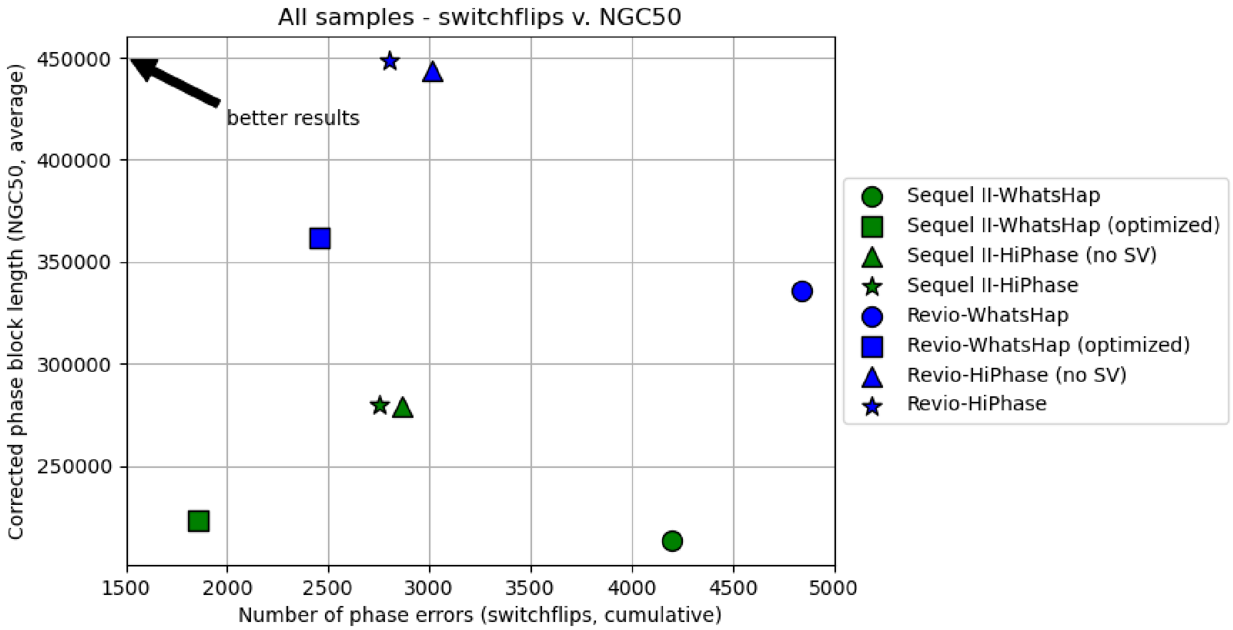

Figure 2: Summary figure showing switchflips v. NGC50 for all conditions ({system}-{tool}). Ideally, there are higher NGC50 values and lower switchflip errors. HiPhase generated fewer switchflip errors than the primary comparator "WhatsHap", but more errors than the "WhatsHap (optimized)" version. However, NGC50 values were consistently higher for HiPhase than either WhatsHap method. Additionally, HiPhase runs on Revio system datasets generated longer NGC50s with comparable error profiles to those from Sequel II systems. In constrast, WhatsHap tended to generate more errors on Revio system datasets than the Sequel II system datasets.

Table 6: Summary metrics for Sequel II system datasets (HG001, HG002, and HG005) for each method. Switchflips, hamming distance, and NGC50 are measures of error. NG50 and NGC50 are measures of phase block length. Number of phased variants is broken into "all" category and the structural variants (SVs) from pbsv. Switchflips, hamming distance, and the number of phased variants are totaled across all datasets (sum), whereas NG50, NGC50, and genes full phased are averaged (mean). Metrics marked with an asterisk (*) are derived from tool-specific sources, details of which are available in the Section 3.1. **Bolded** values represent the best performance in the row.

| **Metric** | **WhatsHap** | **WhatsHap (optimized)** | **HiPhase (no SV)** | **HiPhase** |
|---|---|---|---|---|
| Switchflips | 4,196 | **1,856** | 2,866 | 2,750 |
| Hamming distance | 110,379 | **71,497** | 122,609 | 125,574 |
| NG50* | 248,730 | 238,626 | 303,282 | **306,628** |
| NGC50 | 213,308 | 223,106 | 278,805 | **279,726** |
| Fully phased genes | 88.6% | 86.4% | 94.9% | **95.1%** |
| Number of phased variants (all)* | 8,608,868 | 8,467,279 | 8,852,826 | **8,897,358** |
| Number of phased SVs (pbsv) | N/A | N/A | N/A | **75,869** |

of phased variants, and the location errors. We think of this as a measure of the *severity* of errors in the phase result. In internal testing, we found this metric to be highly variable for an individual dataset. Minor changes in switchflips could lead to much larger changes (both increases and decreases) in the measured hamming distance. In aggregate, it is often useful for assessing trends in error severity. On an otherwise correct phase block with $V$ variants, one flip always increases hamming distance by 1 whereas one switch may increase the hamming distance by up to $\frac{V}{2}$.

Additional details on each can be found here: https://whatshap.readthedocs.io/en/latest/guide.html#whatshap-compare

###### 3.1.2 Phase block metrics

These metrics are used to aggregate phase block statistics. For an individual phase block, the length is measured in base pairs (bp) from the position (POS field in VCF) of the first phased variant to the last phased variant (inclusive):

- NG50 - The smallest phase block length such that all blocks of this size or larger span at least 50% of the full genome (GRCh38) length.
- NGC50 - "Corrected NG50", this metric combines *length* with *accuracy*. Phase blocks are first split wherever a switchflip error occurs and then NG50 is re-calculated on the collection of smaller, error-free sub-blocks. Anecdotally, this metric is useful for defining trade-offs between reducing errors and increasing phase block length (as in "WhatsHap (optimized)" mode).
- Number of phased variants - The number of phased variants across all phase blocks, usually separated by variant type.
- Fully phased genes - The percentage of RefSeq genes where all heterozygous small variants are fully spanned by a single phase block.

##### 3.2 Error metric command templates

Error metrics were gathered in a *homogenized* way by running `whatshap compare` against the corresponding benchmark set. Program 5 is the command template used in our benchmarking pipeline.

Program 5: Command for running `whatshap compare` to gather error metrics on both WhatsHap and HiPhase outputs.

```
whatshap compare \
  --tsv-pairwise {output.tsv} \
  --switch-error-bed {output.error_bed} \
  {input.truth} \
  {input.vcf}
```

Key outputs from this command are:

1. Metrics `switches`, `flips`, and `switchflips` - the column `all_switchflips` from {`output.tsv`} is parsed into separate switch and flip categories and the result for all chromosomes are added together into a single value for the dataset; i.e. `switchflips = switches + flips`
2. Metric `hamming_distance` - the column `blockwise_hamming` from {`output.tsv`} is parsed and the result for all chromosomes are added together into a single value for the dataset
3. The location of each switch error is in {`output.error_bed`}.

##### 3.3 Phase block metric command templates

Metric gathering tools from WhatsHap are limited in that they cannot capture phase results from structural variants or multi-allelic variation. For WhatsHap results, this is not a problem because it does not attempt to solve those variant types. However, HiPhase is capable of including those types in the phase result, so we had to build the results into the HiPhase outputs in order to accurately capture them. The following sections describe how each phase block metric was gathered for each tool.

###### 3.3.1 NG50 and number of phased variants

**WhatsHap** For WhatsHap, we used the built-in `stats` command to gather summary results. Program 6 contains the command template for this tool.

Program 6: Command for running `whatshap compare` to gather phase statistics on WhatsHap phase results.

```
whatshap stats \
  --tsv {output.tsv} \
  --block-list {output.blocks} \
  {input.vcf}
```

Key outputs from this command are:

1. Metrics `NG50` and number of phased variants - for both metrics, we parsed row `ALL` from {`output.tsv`} and collected columns `block_ng50` and `phased`, respectively.
2. The location of each phase block is in {`output.blocks`}.

**HiPhase** For HiPhase, we used the built-in options to gather summary results while phasing (Program 7).

Program 7: Partial command for running `hiphase` and collecting metrics at the same time.

```
hiphase \
  --blocks-file {output.blocks} \
  --summary-file {output.summary} \
  ...
```

Key outputs from this command are:

1. Metrics `NG50` and number of phased variants - for both metrics, we parsed row `all` from {`output.summary`} and collected columns `block_ng50` and `num_phased`, respectively.
2. The location of each phase block is in {`output.blocks`}.

Initially, we were using the same command from Program 6 to gather statistics for HiPhase results. However, WhatsHap does not correctly assess multi-allelic sites, a problem shared in both the `phase` and the `stats` components of the tool. As a result, the `stats` command was consistently *under*-reporting the performance of HiPhase both in the number of variants and the phase block lengths, which led us to develop the custom statistic outputs that are built into HiPhase now. In general, the statistics from Programs 6 and 7 are strongly correlated but more correct with the HiPhase outputs.

###### 3.3.2 NGC50 and "Fully phased genes"

Given the above output phase blocks (which are calculated differently), we *homogenized* the calculation of the NGC50 and the percentage of genes that were fully phased.

1. NGC50
   (a) Per dataset and method, we converted the reported phase block file (TSV/CSV) to a corresponding BED file by extracting the appropriate columns. This is {`input.phase_block_bed`} in Program 8.

(b) The regions overlapping switchflip errors in the {output.error_bed} file, generated during Section 3.2, are removed from the phase block bed file using Program 8. This will split any phase blocks into sub-blocks that contain no errors relatively to the phase benchmark set.

(c) We then re-calculated NG50 using the remaining, phased sub-blocks from {output.correct_phase_block_bed} to get NGC50.

Program 8: Command template for creating correctly phased sub-blocks from a tool's original full phase blocks.

```
bedtools subtract \
  -a {input.phase_block_bed} \
  -b {input.error_bed} > \
  {output.corrected_phase_block_bed}
```

2. "Fully phased genes"

(a) We downloaded the latest RefSeq GRCh38 GFF3 file using this interface: https://www.ncbi.nlm.nih.gov/projects/genome/guide/human/index.shtml. Our downloaded version was annotated with "NCBI Homo sapiens Annotation Release 110".

(b) We performed a one-time conversion of this file into a BED file, keeping only "gene" and "pseudogene" annotations from these sources: "BestRefSeq", "RefSeq", "Gnomon", "Curated Genomic", and "BestRefSeq,Gnomon". Additionally, we only kept annotations on primary chromosomes (all 22 autosomes, chrX, chrY, and chrM). This file is {input.refseq_gene_bed} in Program 9.

(c) Per dataset and method, we converted the reported phase block file (TSV/CSV) to a corresponding BED file by extracting the appropriate columns. We then extended each phase block (both downstream and upstream) until it encountered another heterozygous variant in the dataset's small variant VCF file. These "extended" phase blocks represent the maximum extension of a phase block, intended to capture homozygous regions within genes. This is {input.phase_block_bed} in Program 9.

(d) Per dataset, we ran `bedtools intersect` to capture all RefSeq gene regions that were fully covered by a single extended phase block using Program 9. The percentage fully phased was calculated by counting the number of remaining regions in {output.intersection} and dividing by the original count in {input.refseq_gene_bed} (e.g. `wc -l`). Steps (c) and (d) are captured in a single Python3 script that wraps `bedtools` and performs the calculations.

Program 9: Command template removing genes from a RefSeq BED file that were not fully covered by a single phase block.

```
bedtools intersect \
  -a {input.refseq_gene_bed} \
  -b {input.phase_block_bed} \
  -f 1.0 -wa > \
  {output.intersection}
```

##### 3.3.3 Number of phased structural variants

HiPhase is the only approach that phases structural variants in our tested method. To gather the number of phased structural variants, we initially ran `whatshap stats` as in Program 6, but this produced incorrect results. Instead, we developed a custom Python3 script that parsed the VCF using `cyvcf2` (Pedersen and Quinlan, 2017b), counted variants with the appropriate statuses, and produced a JSON file. An example of the output is shown in Program 10. Note that `pbsv` produces structural variant calls down to approximately 20 bp, so there is some overlap between variants from DeepVariant and `pbsv`. We do not attempt to filter or separate this overlap out in any of our calculations, instead preserving the full DeepVariant and `pbsv` outputs.

Program 10: Example JSON output for structural variant statistics gathering.

```
{
    "heterozygous": 32625,
    "phased": 25960,
    "singletons": 1195,
    "unphased": 6665,
    "variants": 52271
}
```

##### 3.4 Computational resource metrics

We have documented computational resource usage for the tested tools in Table 7. All results were gathered using the built-in `benchmark` option of snakemake (Mölder et al., 2021), details on this functionality can be found at this link: https://snakemake.readthedocs.io/en/stable/tutorial/additional_features.html#benchmarking. For our purposes, we gathered the max memory usage (`max_rss`), wall clock time (`s`), and CPU time (`cpu_time`) for a process. For HiPhase, a single job was used to run each dataset and 16 threads were allocated per job, so those results are simply copied directly from the benchmark outputs and merged into a single value for the method. Unfortunately, WhatsHap does not have built-in parallelization. For efficiency, we split the phasing into separate cluster jobs by chromosome (25 total sub-jobs: 22 for the autosomes plus chrX, chrY, and chrM) and then merged the results. This required us to also merge the compute resources used from benchmarking as follows:

- Wall clock time - *max* of all sub-jobs' wall clock time; if innately parallelized by chromosome, the longest single-chromosome run-time would be the limiting factor
- CPU time - *sum* of all sub-jobs' CPU time; if innately parallelized, the program would still theoretically use the same amount of CPU (if not more from overhead)
- Max memory usage - *max* of all sub-jobs' max memory usage; if innately parallelized, this would likely be higher due to parallel processing and memory consumption (e.g., summation); however, it is difficult to fairly assess this, so we left it as the maximum, which is likely a lower bound for a parallelized WhatsHap

We did not penalize WhatsHap for the overhead of splitting and merging the input and output files, but there was additional compute associated with those steps. Additionally, we note that we ran these analyses in a cluster setting. Readers should be aware that compute times may reflect cluster congestion or other factors that are difficult to isolate in a cluster environment.

#### 4 Deep Methods

**NOTE:** This section is intended to provide deeper information on how HiPhase methods work. Some sections from the main document are repeated here for continuity for the reader.

At a high level, the phasing problem can be broken apart into three major components: phase block generation, allele assignment, and diplotype solving. Phase block generation is the process of generating putative phase blocks by looking for pairs of adjacent heterozygous variant calls that are overlapped by at least one read mapping. These can be chained together to form a candidate phase block, and each one can be processed independently with respect to the next two components. Allele assignment is the process of converting all read mappings (i.e., observations) within a putative phase block into allelic observations, which are chains of reference or alternate alleles corresponding to the observed variants within the particular read mapping. Diplotype solving is the process of distilling these allelic observations into two representative haplotypes that are *expected* to complement each other (i.e., where one is the reference allele, the other is the alternate allele). In the following sections, we describe each of these components in greater detail, focusing on similarities and differences with existing approaches.

Table 7: Compute resources required for each method. Best results in each column are **bolded**. Note that these resources are cumulative (e.g., summation) for all datasets in the corresponding category. Additionally, compute and memory resources were merged separately for WhatsHap due to the lack of innate parallelism. In general, "HiPhase (no SV)" used less compute and wall clock time but more memory than either WhatsHap approach (note: memory consumption may be misleading due to our max memory measure for WhatsHap). Using structural variants and global mode ("HiPhase") increased the compute costs above that of WhatsHap, and further increased max memory consumption as well.

| System | Method | Wall clock time (sum, seconds) | CPU time (sum, seconds) | Max memory usage (sum, GB) |
|---|---|---|---|---|
| Sequel II | WhatsHap | 8,092 | 87,961 | **5.2** |
| | WhatsHap (optimized) | 9,340 | 102,694 | 7.0 |
| | HiPhase (no SV) | **6,292** | **36,411** | 12.7 |
| | HiPhase | 10,531 | 142,362 | 19.5 |
| Revio | WhatsHap | 8,504 | 91,093 | **5.7** |
| | WhatsHap (optimized) | 9,440 | 113,074 | 8.0 |
| | HiPhase (no SV) | **9,018** | **42,374** | 13.9 |
| | HiPhase | 11,183 | 114,901 | 25.7 |

#### 4.1 Phase block generation

Phase block generation is the process of generating putative phase blocks by looking for pairs of adjacent heterozygous variant calls that are overlapped by at least one read mapping. At a high level, a phase block starts by taking the first (or next) available variant on a chromosome and creating a single-variant block (or "singleton" block) from it. Then, the next variant is loaded and HiPhase searches for mappings that span both that new variant and the existing block. If no mappings span both, the algorithm will then check for supplementary mappings from the new variant into the existing block. To our knowledge, this supplementary mapping check is unique to HiPhase, and allows it to span coverage gaps caused by things like homozygous deletions and reference gaps. If at least one spanning (or supplementary) mapping is identified, then the variant is joined to the current phase block and the process is repeated with the next variant. If no spanning (or supplementary) mappings are found, the current block is returned as a putative phase block and a new single-variant block created with the new variant. We note that HiPhase has parameters to adjust the default behavior for phase block generation that is described above.

Each putative phase block acts as an isolated sub-problem in the full solution. Because each putative block is unconnected by the read mappings at their ends, they represent a lower-bound on the number of phase blocks in the final solution (note: HiPhase assumes no phase block overlaps). Most importantly, each sub-problem can be solved independently, allowing for the remaining steps (allele assignment and diplotype solving) to be performed in parallel for each putative phase block. This forms the basis for multi-threading in HiPhase.

#### 4.2 Allele assignment

Allele assignment is the process of converting all read mappings (i.e., observations) within a putative phase block into condensed allelic observations, which are chains of reference or alternate alleles corresponding to the observed alleles within the particular read mapping. The main idea is to simplify a long-read sequence (e.g., >15 kb) down to a smaller set of integer values representing which alleles are present within the read. Typically, these assignments correspond to reference (REF) or alternate (ALT) alleles, but we also allow for ambiguity, unassigned values, and multi-allelic variation (two ALT alleles at one position). Additionally, each observed allele is assigned a "quality" or "weight" indicating the cost to alter or ignore that allele in the diplotype solving process. Program 11 shows a simple example of converting an observation to its condensed representation.

To our knowledge, HiPhase is unique in that it has two modes for allele assignment: local re-alignment and global re-alignment. In brief, local re-alignment assigns alleles and quality based on a small window around each variant position (conceptually similar to the allele assignment process of WhatsHap (Patterson

Program 11: Generic outline of how allele assignment works. An observed sequence (1) is checked for each allele (2-4) and alleles are stored sequentially (5). These are condensed into a final integer representation where only the alleles are left (6).

```
1 Sequence   : ACGAGTTTA
2 Pos 3 A>G  :   G  | |  ALT
3 Pos 6 T>C  :      T |  REF
4 Pos 8 C>G  :        T  AMBIGUOUS
5 Alleles    : --1--0-2-
6 Condensed  : 102
```

et al., 2015)). In contrast, global re-alignment will *fully* re-align the mapping against a local alt-aware reference graph using a graph-aware version of the wavefront algorithm (Marco-Sola et al., 2020). In general, local re-alignment is a faster process, but it is ill-suited for accurate allele assignment for large structural variants and some indels. Global re-alignment tends to be slower but is more accurate when it comes to allele assignment, especially in structural variants. HiPhase implements a "dual mode" allele assignment where if global re-alignment is too slow, it will fall back on local re-alignment for the phase block.

Once all mappings have been converted to a condensed allele representation, HiPhase has one final step where mappings with the same read name are collapsed into a single entry. The primary purpose of this step is to create a bridge between supplementary mappings that span a gap in coverage. This allows HiPhase to cross deletion events and reference gaps with split read mappings covering them. If the mappings for one read overlap but have a conflicting allele assignment, then that allele is converted to an ambiguous allele assignment in the collapsed representation. In the end, each read is represented exactly once in the collection of condensed alleles for the phase block.

##### 4.2.1 Local re-alignment

Local re-alignment is an approach that is very similar to the allele assignment algorithm used by WhatsHap (Patterson et al., 2015). For an individual variant, local re-alignment takes the sequence overlapping the variant with a surrounding window ($\pm W$ base pairs) and performs two alignments to two different sequences: one version that matches allele 0 (typically REF allele) and one version that matches allele 1 (typically ALT allele). Whichever alignment produces the lowest cost (e.g., smallest edit distance) is selected as the allele for that variant. In the event of a tie, they are equidistant, and the allele is marked as ambiguous.

In HiPhase, the default window size is $W = 15$. Additionally, the tool will truncate this window if it overlaps other known variants from the VCF or if it detects unknown variant interference (e.g., a large insertion in the sequence). During development, this tended to produce more accurate allele assignments and reduce ambiguity in the assignments as well.

Once an allele is assigned, it is also given a quality value corresponding to the confidence that the allele is correct. For small variants (SNVs and indels), this quality value is a function of the base quality of the bases in the window that is then scaled depending on the variant type. For example, SNV variants tend to be the most accurate variant calls so they are given increased weight. Insertions, deletions, and indels are all down-weighted with respect to SNVs because they tend to generate more false positive calls. For HiPhase, these static weights were set via heuristics and may benefit from further tuning or some other form of dynamic weighting in the future.

Local-realignment tends to be ill-suited for structural variants. For different mappings, a structural variant can have drastically different coordinates due to local sequence similarity and subtleties in mapping. For window-based approaches, this makes it much easier to miss a true non-reference allele and incorrectly label it as reference allele. While we do not recommend using local re-alignment for phasing structural variants, it is currently supported by HiPhase (partially because of dual mode allele assignment). Structural variant insertions are handled in an identical manner to small insertions using the local window with realignment. Structural variant deletions use an overlap scoring scheme to determine whether an event is present or not, similar to how CNV benchmarking tools like Truvari (English et al., 2022) determine whether a variant call matches. In short, it counts the number of deleted bases in the mapping that overlap the deletion call. If a sufficient number of bases are marked as deleted in the mapping, then it will assign it the deletion allele. The quality of this event then scales based on how well it overlaps (e.g., if there is a 70% match, it will receive 70% of the maximum quality). Given the inaccuracy of structural variant local re-alignment, these events are down-weighted relative to other variant types.

###### 4.2.2 Global re-alignment

To our knowledge, the application of global re-alignment to phasing is novel and unique to HiPhase. In contrast to local re-alignment, this approach attempts to fully re-align the entire read mapping against an ALT-aware graph sequence that is localized to the mapping. Using the local reference genome as a backbone for the graph, all provided variants (including homozygous variants that are not part of the phasing problem) are added sequentially to the graph structure. This approach allows methods like Partial-Order Alignment (POA) (Lee et al., 2002) to run on top of the graph. The core idea is to find the lowest cost path through the graph while also tracking the nodes (which correspond to alleles) that were traversed in that path. For those unfamiliar with POA, the Simpson Lab provides an intuitive explanation for those who are already familiar with pairwise alignment algorithms: https://simpsonlab.github.io/2015/05/01/understanding-poa/.

Unfortunately, the original POA approach can be quite slow on reads that are >10 kb in length due to scaling off of both the graph size and mapping length ($O(G * N)$ where $G$ is the graph size and $N$ is the mapping length). Additionally, we know that the graph and the mapping *should* be very similar since the read mapped to that location, a property that is not taken advantage of in the original POA approach. In contrast, the pairwise WFA algorithm (Marco-Sola et al., 2020) is specifically designed to leverage sequence similarity between two sequences, with a run-time of $O(N * s)$ where $s$ is the number of differences between the two sequences. However, this algorithm was not designed to run on a graph structure.

HiPhase combines the benefits of POA and WFA into a novel implementation of the WFA algorithm that is designed to run on a localized reference graph structure. This algorithm leverages the benefits of WFA (specifically, scaling off of the number of differences, for run time of $O(G * s)$), while simultaneously tracking the optimal series of nodes that were traversed in the graph. The result is an efficient, lowest-cost traversal of the localized graph that identifies the optimal allele assignments for the read mapping. We refer to this method as "graph WFA" in subsequent sections.

In theory, one could perform a full backtrace of the graph WFA algorithm to get base-level quality values, but this can be expensive and tricky to handle when ambiguity in node traversal is present in the re-alignment (e.g., an indel assignment is truly ambiguous, but different lengths of read sequence match the two options). Instead, global re-alignment in HiPhase uses *only* the variant type to determine allele quality scores. In general, the relative weights used by global re-alignment match those from local re-alignment (e.g., SNV has the highest weight, while indels are down-weighted relative to SNVs). As noted for local re-alignment, these static weights were set via heuristics and may benefit from further tuning or some other form of dynamic weighting in the future.

While this algorithm is generally fast for most phase blocks, it can still run prohibitively long in noisy areas or regions with lower accuracy in variant calling (these both increased the error term, $s$, above). To prevent excessive run-times, HiPhase enforces a per-block, CPU-time limit for global re-alignment. In the event that the user-provided CPU-time is exceeded for a given phase block, any work so far will be discarded, and the algorithm will revert to local re-alignment for *all* mappings in the block. This ensures that all allele and quality assignments for a given block use the same approach. In our test cases, the vast majority (>99.8%, see Table 9) of phase blocks succeed in global re-alignment.

We note that while the HiPhase implementation of a graph-based WFA approach was done in parallel to and without knowledge of GWFA, the core algorithms converged on very similar solutions. While the application and implementation of the algorithm is slightly different, we recommend reviewing the source code (https://github.com/lh3/gwfa) and the virtual seminar by author Heng Li (https://www.youtube.com/watch?v=um_q8-BOXpg) for greater details on core concepts for graph WFA.

###### 4.2.3 Collapsing mappings

Once all mappings have been converted to their corresponding condensed allele representation, HiPhase has one final step where mappings with the same read name are collapsed into a single entry. The primary purpose of this step is to create a bridge between supplementary mappings that span a gap in coverage. This allows HiPhase to cross many homozygous deletion events and reference gaps despite lacking a single mapping that directly spans the gap. If conflicting alleles are encountered during this process (i.e., the mappings overlap and have a different assigned allele), then the allele is converted to an ambiguous representation. Figure 3 shows an example where this process allows HiPhase to create a single phase block spanning a reference gap where WhatsHap created two separate blocks. At the end of the collapsing process, each read is represented exactly once in the collection of condensed alleles for the phase block.

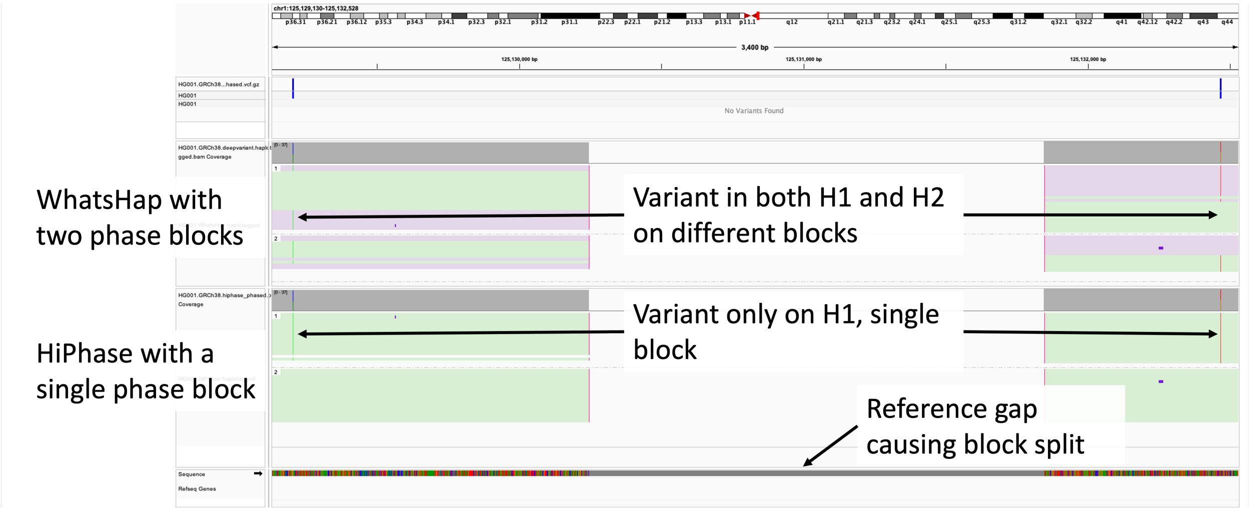

Figure 3: IGV screenshot of a reference gap in HG001. The top haplotagged BAM track is from WhatsHap, and the bottom haplotagged BAM track is from HiPhase. Mappings in both tracks are grouped by haplotag ID (HP:1 or HP:2) and colored by phase set (PS). In the WhatsHap track, there are two blocks corresponding to upstream (green) and downstream (purple) phase blocks. Read mappings have supplementary mappings across the gap but are assigned to different blocks and haplotypes. Additionally, heterozygous ALT variants immediately upstream and downstream do not cleanly segregate into only one haplotype, appearing in both HP:1 and HP:2 depending on the assigned block ID. In contrast, HiPhase has a single phase block (green) spanning the region, and both heterozygous ALT alleles are cleanly assigned to HP:1.

#### 4.3 Diplotype Solving

Diplotype solving is the process of distilling the condensed allelic observations into two representative haplotypes that are *expected* to complement each other (i.e., where one is the reference allele, the other is the alternate allele). For our purposes, we define the core diplotype solving problem as a slight reformulation of the weighted minimum error correction (wMEC) problem as described by the authors of WhatsHap (Patterson et al., 2015). Given a matrix where each row corresponds to the condensed allele representation of one read in the phase block, the goal is to find two haplotypes, $h_1$ and $h_2$, such that the cost of changing each row to exactly match *either* $h_1$ or $h_2$ is minimized. Additionally, each allele has a cost or "weight" associated with altering that allele. We refer the reader to (Patterson et al., 2015) for greater technical details around this and other formulations of the phasing / wMEC problem.

##### 4.3.1 A* phasing algorithm

HiPhase uses a version of the A* search algorithm (Hart et al., 1968) to solve the phasing problem. In general, A* search algorithms explore some search space by iteratively expanding the current lowest cost option (similar to Dijkstra's algorithm (Dijkstra, 2022)). The key difference with A* is that cost for a partial solution or "node" $n$ is defined by both an observed cost, $g(n)$, and a heuristic estimated cost to reach the final goal, $h(n)$. As long as the heuristic is "admissible", meaning it never *over*-estimates the cost to the goal, A* is guaranteed to find the optimal solution. Thus, the key challenges in all A* search algorithms are 1) defining the search space, 2) defining the cost structure, and 3) defining an efficient, admissible heuristic that is *ideally* close to the true cost. We break down each of these components for HiPhase in the following sections.

##### 4.3.2 Defining the search space

Given our wMEC problem formulation, the search space is relatively simple: we are searching through all possible combinations of $h_1$ and $h_2$. Conceptually, this can be visualized as a search tree where the root contains empty haplotypes (i.e., $h_1 = h_2 =$ “”). All unexpanded nodes (initially just the root) are added to a priority queue that is based on the lowest cost (defined by $g(n) + h(n)$). In each iteration, the lowest cost node is removed from this queue and expanded. This expansion process adds one new node for each possible extension of the partial solution represented by the current lowest code node. In the expected case, this would be one of two phase orientations: 0|1 and 1|0. However, we also allow for variants to be converted to homozygous if that leads to a more optimal solution: 0/0 or 1/1 (in output files, these are left as unphased heterozygous calls, 0/1). This leads to a total of four expansions per node (two heterozygous and two homozygous). Given these expansion definitions and the initial empty root node, we can also say that each node at depth $D$ contains only haplotype combinations of length $D$. Thus, for a phase block of length $N$, this tree has a *maximum* depth of $N$ and has at most $O(4^N)$ nodes in a fully expanded tree. Finally, the first node at some depth, $d$, that is popped from the priority queue represents the optimal solution for the first $d$ variants in the block. This is because all other nodes have a cost that is greater than or equal to this node, and due to the admissibility constraint of $h(n)$, we know that the costs of those nodes when extended can only go up. Thus, the first node in the traversal that is encountered at depth $N$ is the solution to the full phase block.

In practice, we found that the majority of the phase blocks have very clean solutions, leading to near linear tree traversals (e.g., $O(N)$). However, some phase blocks have less clean solutions that create less direct, sometimes even exponential, tree traversals. These blocks tend to be clustered around problematic genomic regions such as segmental duplications, low complexity regions, centromeres, or other high-mismatch regions. If left unchecked, they would likely have near-exponential run-times (e.g., $O(4^N)$) to reach the true optimal solution. To resolve this issue, we implemented a pruning strategy that limits the number of nodes in the priority queue. When the limit is reach, a pruning threshold, $p$, is incremented and all nodes that are not at least $p$ deep in the tree are pruned. Functionally, this is pruning nodes that are relatively shallow in the tree exploration and less *likely* to contain the optimal solution. However, if *anything* is pruned this way, A* loses the guarantee of finding the optimal solution as the partial solution that was pruned may eventually lead to the true optimum. HiPhase tracks phase blocks where pruning occurs which may be useful for assessing phase block quality by users or downstream tools. On our test data, HiPhase has unpruned, guaranteed optimal solutions for approximately 88-93% of phase blocks depending on the dataset and allele assignment mode (see Table 9).

##### 4.3.3 Defining the cost structure

When defining the cost of a given pair of partial haplotypes, we must compare those partial haplotypes to our condensed allele observations and the corresponding weights assigned to each allele. For a node $n$ at depth $d$, we have two partial haplotypes of length $d$, $h_{n1}$ and $h_{n2}$, that form partial candidate solutions for phasing the full block. Each read observation in the phase block is compared to the two haplotypes and assigned a cost of converting that read to have no conflicts with the haplotype. Then, the minimum of these two values is selected as the cost for this read and the node, effectively assigning it to one of the two partial haplotypes. The sum of all costs represents the total observed cost, $g(n)$, for node $n$. Mathematically, given a collection of reads, $R$, where each read has both alleles and weights, this can be represented with two formulas. The first formula (Equation 1) defines the cost function for an individual comparison of a read $r$ to a partial haplotype $h$, and the second formula (Equation 2) defines the combined cost across the read collection when compared to both partial haplotypes associated with node $n$:

$$\text{cost}(r, h) = \sum_{i=0}^{|h|} \begin{cases} 0, & \text{if } r.allele[i] = h[i], \\ r.weight[i], & \text{otherwise} \end{cases} \tag{1}$$

Table 8: Table showing a simplified version of how the heuristic chain is built to create heuristic estimated costs for all nodes at depth $d$. Sub-problem solutions are initial generated, in this case subproblems of length 2 (e.g. two variants phased together), and stored as $P$. Then the heuristic chain is constructed in reverse order. The last 2 nodes simply copy the subproblem solutions. Then, starting with variant $C$, the chaining process looks at both the subproblem solution and the chain solution that is 2 away ($h(E)$ for variant $C$). This process continues in reverse order until the full heuristic is solved starting at variant $A$. In practice, this forms a monotonically decreasing array for the heuristic costs, $h(d)$.

| **Variant index,** $v$ | **A** | **B** | **C** | **D** | **E** |
|---|---|---|---|---|---|
| **Subproblem cost,** $P_v$ | 3 | 2 | 5 | 4 | 0 |
| **Heuristic cost,** $h(d)$ | $h(A) = P_A + h(C) = 3 + 5 = 8$ | $h(B) = P_B + h(D) = 2 + 4 = 6$ | $h(C) = P_C + h(E) = 5 + 0 = 5$ | 4 | 0 |

$$g(n) = \sum_{r}^{R} \min(\text{cost}(r, h_{n2}), \text{cost}(r, h_{n1})) \quad (2)$$

While not specified in the above formula, all weights for ambiguous or unassigned alleles are set to 0, so there is no cost associated with reads that have unassigned alleles because they do not fully span a phase block. This means that once a node's haplotypes have extended past the last set allele for a read, the cost of that read becomes fixed for all extensions of that node (e.g., there is no weight on the read past that point that may change the minimum cost). In practice, this allows HiPhase to separate the cost of $g(n)$ into costs associated with "frozen" and "liquid" reads. Frozen reads will *never* change cost value as the haplotypes are extended, whereas liquid reads may change by the lowest cost flipping from one haplotype to the other. Frozen read costs are stored in aggregate and not recomputed with each extension to reduce compute time.

###### 4.3.4 Defining the heuristic

Given the above definitions, the A* algorithm could run by simply setting the heuristic component to zero, $h(n) = 0$, which functionally leads to Dijkstra's algorithm (Dijkstra, 2022). However, this would be a very poor heuristic leading to over-traversal of the tree. Ideally, there is a heuristic that is very close to the actual final cost of the solution. While not guaranteed, heuristics closer to the actual cost *tend* to reduce run-time for most A* algorithms.

In HiPhase, we calculate the heuristic by solving sub-problems from the full block. Conceptually, if a phase block has $N$ variants to phase, the block can be broken into sub-problems of some length, $S \leq N$. Any solution to the full phase block *must* also span the sub-problems as well. Let $P_0$ represent the first sub-problem and $P_{N-S}$ be the last sub-problem and assume *some* algorithm can find an optimal solution to these sub-problems. One way to estimate the *minimum* cost of the full solution is to find a long chain of non-overlapping sub-problems from $P_0$ to $P_{N-S}$. The optimal solutions from these sub-problems can then be added together to form a full estimate. While these optimal sub-problem solutions may not be part of the final solution (they are locally optimal, not necessarily globally optimal), any solution to the full problem *cannot* create a solution with less cost than the chain of locally optimal subproblems (it may be equal). Additionally, this property is not limited to the full phase block but can be applied to any partial solution as well. For example, if we wanted to estimate the cost from $P_q$ to $P_{N-S}$, we could run a similar chaining approach starting from $P_q$ instead of $P_0$. This means that for any point in the full phase block, we have a general strategy to estimate the distance to the end of the block, which we use for our heuristic estimate, $h(n)$. In Table 8, we show a simplified example of the full heuristic being constructed for a toy problem.

There are several subtleties to how HiPhase does this in practice. First, given the above strategy, the heuristic estimates are tied to the variant index in the phase block, which is also the depth in the search tree (i.e., $h(n) = h(\text{depth of } n) = h(d)$). This means all nodes at a given depth, $d$, will have the same heuristic estimate to reach the end of the phase block. Second, HiPhase calculates the heuristic in reverse-linear order such that $P_{N-S}$ is the first solved sub-problem and $P_0$ is the last. This allows HiPhase to build up the full heuristic chain as it goes (e.g., $h(q) = h(q + S) + P_q$). Third, while we focus on fixed-size sub-problems in our description, HiPhase will compute all sub-problems of size $\leq S$. This allows for chaining a different number of sub-problem solutions of potentially different sizes (e.g., 10 variants could be sub-problems of size $(5 + 5)$,

$(3 + 3 + 4)$, etc.). Fourth, while the sub-problem solver is looking for *locally minimal* cost solutions, the heuristic is looking for the *maximum* cost chain of these minimal solutions. Intuitively, this is because if some chain of minimal sub-problem solutions exists with some cost, we know that any full phase block solution *must* have at least that cost and likely more from joining the full chain of sub-problem solutions together. Thus, it looks for the most costly chain of non-overlapping sub-problems to obtain the closest estimate to the actual cost. Finally, HiPhase uses a recursive A* phasing algorithm to solve the sub-problems. This recursive approach is generally identical to the full unpruned A* phasing algorithm, but it is limited to a fixed number of node expansions to reduce the computational burden of solving each sub-problem. This means that the heuristic will not always use the full sub-problem solution of length $S$, but may instead terminate early.

#### 4.4 Algorithm Statistics

HiPhase can output statistics about each block and the performance of the underlying algorithms on that block. Table 9 shows some summary statistics on the allele assignment and A* algorithm execution for the datasets used in our results. Each statistic is briefly described here:

- Percent blocks globally re-aligned – When global re-alignment is enabled, this is the percentage of putative phase blocks that successfully finished the global re-alignment process within the allotted CPU time.
- {`variant_type`} REF:ALT ratio – For a given {`variant_type`}, this is the ratio of alleles that were assigned to the reference (REF) or alternate (ALT) allele. Ideally, this ratio is near 50:50, indicating an unbiased balance in allele assignment. Deviations from 50:50 may indicate errors in allele assignment and/or errors in upstream variant calling (e.g., false positives or inaccurate positions / ALT sequences).
- Heuristic / actual cost – Across all blocks, this is the total heuristic estimated cost divided by the actual cost in the final solutions. A perfect heuristic has a value of 1.0, indicating that it *exactly* estimated the actual cost. While not a guarantee, as this value approaches 1.0, the A* algorithm *tends* to converge on a solution faster by exploring fewer nodes.
- Unpruned, exact solutions – The percentage of blocks that converged on a solution without performing any pruning of the search space. From an A* phasing perspective, these solutions are *guaranteed* to be optimal given the problem design. Note that optimal does *not* always mean biologically correct, but we expect these to be correlated given the problem design.

Table 9 includes a comparison of three modes of running HiPhase. "Small variants only" is the same as "HiPhase (no SV)" and "Small and structural variants (global, 300 sec)" is the same as "HiPhase" in the main manuscript. The "local" mode phases both small and structural variants, but without global re-alignment enabled. As noted in Section 4.2.1, we do not recommend this mode for assigning structural variant alleles but include it here for demonstrating the benefits of global re-alignment mode.

Table 9 highlights a few notable differences between local and global modes. First, with global re-alignment, the ratio of alleles assigned to the ALT haplotypes improves for both structural variant deletions and structural variant insertions, with the greatest improvement in insertions. This is most likely because the global re-alignment process is more capable of handling noise in the mapping location of large insertions. Second, the number of exact solutions generated while using global mode increased slightly ($\sim$1%), indicating that the algorithm can more easily phase the blocks as a result of more accurate allele assignments. Finally, while global re-alignment can timeout in difficult regions, the vast majority of the phase blocks succeeded (>99.8%) within the given time limit.

Table 9: Algorithmic performance of the HiPhase implementation of global re-alignment and A* phasing on Sequel II system and Revio system datasets. The first three statistics are gathered from the allele assignment process, and the last two are from the A* phasing algorithm. Structural variant (SV) deletion and insertion ratios are separated to show the separate impact on each type. The best metric for each row is **bolded**.

| **System** | **Metric** | **Small variants only (local)** | **Small and structural variants (local)** | **Small and structural variants (global, 300 sec)** |
|---|---|---|---|---|
| Sequel II | Percent blocks globally re-aligned | N/A | N/A | **99.88%** |
| | SV Deletion REF:ALT ratio | N/A | 57.93 : 42.07 | **55.48 : 44.52** |
| | SV Insertion REF:ALT ratio | N/A | 71.89 : 28.11 | **61.15 : 38.85** |
| | Heuristic / actual cost | **0.8761** | 0.8711 | 0.8327 |
| | Unpruned, exact solutions | 93.08% | 92.36% | **93.65%** |
| Revio | Percent blocks globally re-aligned | N/A | N/A | **99.81%** |
| | SV Deletion REF:ALT ratio | N/A | 58.54 : 41.46 | **56.55 : 43.45** |
| | SV Insertion REF:ALT ratio | N/A | 73.85 : 26.15 | **67.22 : 32.78** |
| | Heuristic / actual cost | **0.8830** | 0.8791 | 0.8400 |
| | Unpruned, exact solutions | 89.32% | 88.62% | **90.39%** |
